# Loss of *Bacteroides thetaiotaomicron* bile acid altering enzymes impact bacterial fitness and the global metabolic transcriptome

**DOI:** 10.1101/2023.06.27.546749

**Authors:** Arthur S. McMillan, Matthew H. Foley, Caroline E. Perkins, Casey M. Theriot

## Abstract

*Bacteroides thetaiotaomicron* (*B. theta*) is a Gram-negative gut bacterium that encodes enzymes that alter the bile acid pool in the gut. Primary bile acids are synthesized by the host liver and are modified by gut bacteria. *B. theta* encodes two bile salt hydrolases (BSHs), as well as a hydroxysteroid dehydrogenase (HSDH). We hypothesize that *B. theta* modifies the bile acid pool in the gut to provide a fitness advantage for itself. To investigate each gene’s role, different combinations of genes encoding bile acid altering enzymes (*bshA, bshB*, and *hsdhA*) were knocked out by allelic exchange, including a triple KO. Bacterial growth and membrane integrity assays were done in the presence and absence of bile acids. To explore if *B. theta’s* response to nutrient limitation changes due to the presence of bile acid altering enzymes, RNASeq analysis of WT and triple KO strains in the presence and absence of bile acids was done. WT *B. theta* is more sensitive to deconjugated bile acids (CA, CDCA, and DCA) compared to the triple KO, which also decreased membrane integrity. The presence of *bshB* is detrimental to growth in conjugated forms of CDCA and DCA. RNA-Seq analysis also showed bile acid exposure impacts multiple metabolic pathways in *B. theta*, but DCA significantly increases expression of many genes in carbohydrate metabolism, specifically those in polysaccharide utilization loci or PULs, in nutrient limited conditions. This study suggests that bile acids *B. theta* encounters in the gut may signal the bacteria to increase or decrease its utilization of carbohydrates. Further study looking at the interactions between bacteria, bile acids, and the host may inform rationally designed probiotics and diets to ameliorate inflammation and disease.

**Importance:** Recent work on BSHs in Gram-negative bacteria, such as *Bacteroides*, has primarily focused on how they can impact host physiology. However, the benefits bile acid metabolism confers to the bacterium that performs it is not well understood. In this study we set out to define if and how *B. theta* uses its BSHs and HSDH to modify bile acids to provide a fitness advantage for itself *in vitro* and *in vivo*. Genes encoding bile acid altering enzymes were able to impact how *B. theta* responds to nutrient limitation in the presence of bile acids, specifically carbohydrate metabolism, affecting many polysaccharide utilization loci (PULs). This suggests that *B. theta* may be able to shift its metabolism, specifically its ability to target different complex glycans including host mucin, when it comes into contact with specific bile acids in the gut. This work will aid in our understanding of how to rationally manipulate the bile acid pool and the microbiota to exploit carbohydrate metabolism in the context of inflammation and other GI diseases.

## Introduction

The microbiota of the intestinal tract is made up of a diversity of microbes that interact with the host and each other. Bile acid metabolism represents an important function that mediates the relationship between the host and the gut microbiota [1-4]. Bile acids are synthesized from cholesterol in the liver, stored in the gallbladder, and secreted when the host eats a meal [5]. Their role during digestion is to act as surfactants allowing for the absorption of dietary fats and vitamins. Once they reach the terminal ileum, ∼95% of bile acids are reabsorbed through enterohepatic circulation [6, 7]. Bile acids directly interact with the host through receptors like Farnesoid X Receptor (FXR) and Pregnane X Receptor (PXR) that regulate of the production of bile acids made by the host [1, 8], but can also influence other aspects of host physiology, such as immunity by modulating T_reg_ and T_H_17 cell differentiation [8, 9].

Bile acids also shape the gut microbiome’s composition due to their antimicrobial and detergent-like properties [4]. Distinct changes to the bile acid pool and microbiome are associated with a wide variety of diseases including *Clostridioides difficile* infection (CDI) [10], inflammatory bowel disease [11], metabolic syndrome [12], and cancer [12, 13]. However, the mechanisms that dictate the bacterial-bile acid relationship is not fully understood. There is great interest to define these mechanisms in order to leverage the gut microbiota’s ability to alter the bile acid pool, promoting host health.

During their intestinal transit, bile acids are modified by gut bacteria which changes the chemistry and diversity of the bile acid pool. The gut microbiota encodes a variety of bile acid altering enzymes that act on either the sterol core or the amino acid conjugated to it. Bile salt hydrolases (BSHs) are traditionally known to cleave the glycine or taurine amino acid from conjugated bile acids, yet recent work has prescribed a new function for BSHs: the production of bile acids conjugated with a variety of additional amino acids, collectively referred to as microbial conjugated bile acids (MCBAs) [14-16]. Historically, it was proposed that BSHs primary role in bacteria was to detoxify conjugated bile acids thereby promoting bacterial colonization in the harsh gut environment [17]. It has also been hypothesized that BSHs are able to provide nutrients in the form of the liberated amino acid (either taurine or glycine) for the microbe that encodes them or the surrounding gut microbiota [18]. However, a recent study using Gram-positive lactobacilli found that some bile acids became more toxic to the bacterium after deconjugation [19]. The bile acid chemical structure and the different BSH substrate preferences together were able to influence *in vitro* and *in vivo* growth of the different lactobacilli strains [19]. Hydroxysteroid dehydrogenases (HSDHs) were also thought to detoxify bile acids as well as use bile acids as an electron donor for the electron transport chain [20, 21], but this enzyme has been studied even less.

Recent work on BSHs in Gram-negative bacteria, such as *Bacteroides*, has primarily focused on how they can impact host physiology [22, 23]. Bile acid metabolism has been implicated in successful fecal microbiota transplants, cholestatic liver injury, diabetes in bariatric surgery recipients, as well as in a member of the *Bacteroidales* ability to ameliorate hepatic fibrosis [24-27]. However, the benefits bile acid metabolism confers to the bacterium that performs it is not well understood. *Bacteroides thetaiotaomicron* (*B. theta*) is a Gram-negative organism, which encodes multiple genes encoding bile acid altering enzymes [28]. This organism is widely studied due to its unique carbohydrate utilization ability and for this reason has many genetic tools [29, 30]. It is also a prominent member of the human intestinal tract [31]. *B. theta* encodes two putative BSHs (BT_1259 referred to in this paper as BSHa, and BT_2086 referred to in this paper as BSHb) [28]. The majority of bile acid deconjugation in *B. theta* is done by BSHb [23]. Although, there is evidence that BSHa may act on high concentrations of TCA and GCA [32]. The BSHs encoded by *B. theta* have a limited capacity to produce MCBAs thus far, but this requires further study [33]. *B. theta* also encodes a 7α-hydroxysteroid dehydrogenase or HSDH (BT_1911 referred to in this paper as HSDHa) that has not been well studied. αHSDHs act on the sterol core of bile acids to convert the hydroxyl to a ketone that can, in combination with other enzymes, result in the epimerization of hydroxyl groups or remove them entirely [34]. The HSDH encoded by *B. theta* has activity on two major bile acids which contain a 7-hydroxyl group by oxidizing the 7-hydroxyl to a ketone (ex. converting CDCA to 7- oxo LCA) [35].

In this study we set out to define if and how *B. theta* uses its BSHs and HSDH to modify bile acids to provide a fitness advantage for itself *in vitro* and *in vivo*. To address this we created gene knockouts, single and multiple combinations of both BSHs (*bshA* and *bshB*) and HSDH (*hsdhA*) in *B. theta.* We found that *B. theta* could grow in higher concentrations of taurine-and glycine-conjugated CA compared to CDCA and DCA. It was also more sensitive to deconjugated bile acids in growth and in membrane integrity assays. *bshB* was detrimental to *B. theta* growth in the presence of conjugated bile acids that it could act on. RNAseq analysis was done on WT and triple KO strains in rich and minimal media in the presence and absence of bile acids to decipher the relationship between these enzymes and how they affect the global metabolic transcriptome. Genes encoding bile acid altering enzymes were able to impact how *B. theta* responds to nutrient limitation in the presence of bile acids, specifically carbohydrate metabolism, affecting many polysaccharide utilization loci (PULs). This suggests that *B. theta* may be able to shift its metabolism, specifically its ability to target different complex glycans including host mucin, when it comes into contact with specific bile acids in the gut. Bacterial fitness of the WT and triple KO strains showed modest differences *in vivo*, in different mouse models that varied in their bile acid pool. This work will aid in our understanding of how to rationally manipulate the bile acid pool and the microbiota to exploit carbohydrate metabolism in the context of inflammation and other GI diseases.

## Methods

### Bacterial strains and culture conditions

The background for mutant strains is *B. thetaiotaomicron* VPI-5482 Δ*tdk* (WT) that was acquired from the Martens lab at the University of Michigan. *B. theta* strains were grown statically at 37°C in a Coy anaerobic chamber (2.5 % H_2_ /10 % CO_2_ /88.5 % N_2_) in TYG medium (10 g/L tryptone, 5 g/L yeast extract, 4 g/L D-glucose, 100 mM KH_2_PO4, 8.5 mM [NH_2_]_4_SO_4_, 15 mM NaCl, 5.8 μM vitamin K3, 1.44 μM FeSO4⋅7H_2_O, 1 mM MgCl_2_, 1.9 μM hematin, 0.2 mM L-histidine, 3.69 nM vitamin B12, 208 μM L-cysteine, and 7.2 μM CaCl_2_⋅2H_2_O). Solid media for *B. theta* was Brain-heart infusion (Difco) agar supplemented with 10% Horse Blood (LAMPIRE) (BHI-HB). Liquid *E. coli* cultures were grown at 37°C aerobically with shaking in LB (Sigma). Solid media for *E. coli* cultures were grown statically at 37°C aerobically on LB agar. Selective drug concentrations were consistent across solid and liquid media and supplemented during appropriate selection steps (150 μg/mL ampicillin, 200 μg/mL gentamicin, 25 μg/mL erythromycin, 200 μg/mL FUDR)

### Deletion of bsh and hsdh genes from B. thetaiotaomicron

Clean in-frame deletions of bile altering enzymes were generated using a counterselectable allelic exchange procedure [30]. Briefly, 750-basepair regions flanking each gene were amplified by PCR and Hi-Fi DNA Assembly (NEB) was used to clone into pExchange-*tdk*. Plasmids were verified using Sanger sequencing and electroporated into *E. coli* S17 and selected for on solid media supplemented with ampicillin to function as a conjugal donor. An approximately equivalent donor to recipient ratio was used for conjugation. Merodiploids were selected after conjugation on solid *B. theta* media supplemented with gentamicin and erythromycin. A pool of merodiploids was grown in liquid media in the absence of antibiotic to allow for excision of the plasmid from the genome. Successful transformants were selected for on solid *B. theta* media containing FUDR to select against colonies that still contain the plasmid. Single colonies were then verified as successful knockouts using PCR and Sanger sequencing.

### Minimum inhibitory concentrations

BA tolerance measured by MIC was adapted from a previously established bile tolerance assay[19, 36]. Overnight cultures (∼10^10^ CFUs/mL) were inoculated at approximately 10^8^ CFUs/mL into TYG containing a range of BA concentrations. Cultures were anaerobically incubated for a total of 24 h at 37 °C. At both 12 h and 24 h, cultures were serially diluted in PBS and plated on BHI-HB to determine if the concentration of BA tested inhibited growth to prevent at least one doubling of colonies after either 12 or 24 h of growth.

### Propidium iodide staining

Overnight cultures (∼10^10^ CFUs/mL) were diluted to an Opitcal Density or OD_600_ of 0.1 and incubated until an OD_600_ of 0.8 was reached. These concentrations: 4 mM TCA, 4 mM GCA, 5 mM CA, 1.25 mM TCDCA, 1.25 mM GCDCA, 0.3125 mM CDCA, 0.6375 mM TDCA, 0.6375 mM GDCA, 0.3125 mM DCA, or 150 μM SDS were then introduced and incubated at 37 °C for 30 min. Following this incubation bacteria were stained with 20 μM PI using previously described methods [10, 11]. Fluorescence of PI was measured using a TECAN INFINITE F200 (excitation = 560 nm, emission = 600 nm) and was normalized to the OD_600_ of the culture prior to introduction of bile acids.

### *B. theta* tolerance to sub-inhibitory concentrations of bile acids

Overnight cultures (∼10^10^ CFUs/mL) were inoculated 1% into TYG containing a high yet survivable concentration of bile acid that results in approximately 10^10^ CFUs/mL after 12 h (4 mM TCA, 4 mM GCA, 5 mM CA, 1.25 mM TCDCA, 1.25 mM GCDCA, 0.3125 mM CDCA, 0.6375 mM TDCA, 0.6375 mM GDCA, and 0.3125 mM DCA). Cultures were anaerobically incubated for 24 h at 37 °C. At both 12 h and 24 h, cultures were serially diluted in PBS and plated on BHI-HB to determine the effect each bile acid had on the fitness of *B. theta* with and without bile altering enzyme(s).

### RNA extraction from liquid *B. theta* cultures

RNA extraction protocol was adapted from a previously optimized workflow [37]. Overnight cultures were inoculated 1% into TYG or MM supplemented with 4 mM TCA, 4 mM GCA, 2.5 mM CA, 1.25 mM TCDCA, 1.25 mM GCDCA, 0.3125 mM CDCA, 0.637 5mM TDCA, 0.6375 mM GDCA, and 0.156 mM DCA and allowed to reach mid-log (∼1.6 OD) and were fixed 1:1 in a 50/50 Ethanol/Acetone solution and frozen at −80°C. Fixed cultures were thawed on ice and the pellet was resuspended in 1:100 beta-mercaptoethanol. Pellets were then resuspended in TRIzol Reagent (Thermo) and incubated for 20 min. Chloroform was then added and vigorously inverted before incubation at room temperature for 20 min. The samples were centrifuged at 14,000 rpm at 4 °C for 20 min. The aqueous phase (∼650 μl) was then added to 650 μl of isopropanol that had been supplemented with 5 μg/mL glycogen. Samples were vortexed and incubated on ice for 20 min, and then were centrifuged at 4 °C for 30 min. Pellets were washed three times with 70% ethanol, and then dissolved in sterile deionized water. The RNA was treated with Turbo DNase (Thermo Fisher, AM2239); the protocol was modified by increasing the amount of enzyme to 5 μl per sample. After 30 min of incubation in a heat block at 37 °C, 2 μl of Turbo DNase enzyme was added to each sample for a further 30 min of incubation. The RNA was then column purified according to the manufacturer’s instructions (Zymo). Final concentrations of RNA were determined via Qubit™ RNA high sensitivity assay kit.

### Quantitative reverse transcription PCR

RNA from liquid cultures of *B. theta* WT was used as template in reverse transcription reactions using the Murine Moloney Leukemia Virus Reverse Transcriptase (Thermo) following the manufacturer’s protocol. The resulting cDNA was diluted in deionized water such that approximately 10ng would be used as template for quantitative PCR with the SsoAdvanced Universal SYBR Green Supermix (Bio-Rad). Each gene/bile acid combination assayed was analyzed using the ΔΔCt method by comparison to the housekeeping gene 16S rRNA and a control culture grown without bile acid [38].

### RNAseq analysis

rRNA depletion and RNA Sequencing using SP lanes on an Illumina NovaSeq 6000 was performed by the Roy J. Carver Biotechnology Center at the University of Illinois at Urbana-Champaign. Reads were mapped to *B. thetaiotaomicron* VPI-5482 (ASM1106v1) using Salmon with default settings. Differential expression analysis was performed in R using the DESeq2 package. Genes with less than 10 total reads were filtered. For comparisons between WT and triple KO strains of *B. theta* the effect of the nutrient limitation as well as the change in the effect of nutrient limitation due to the genotype was analyzed via the DESeq2 package. For comparisons in growth of bile acids the effect of the addition of bile acid as well as the change in effect of bile acid stress due to nutrient limitation was analyzed via the DESeq2 package. Kegg annotations were downloaded direction from KEGG database and further annotation utilized BLAST KOALA of translated nucleotide sequences to determine K numbers for annotation of unannotated genes. Target substrates and gene annotation of PULs were annotated using CAZy’s PULDB (http://www.cazy.org/PULDB/) and from multiple publications for deeper annotation [39-46].

### Protein cloning and expression

Codon-optimized BT_1911 and BT_1259 were synthesized from IDT and amplified by PCR with the Phusion Flash High-Fidelity PCR master mix (Thermo) using the custom oligonucleotide primers (IDT) listed in the primer table. Amplicons were purified using the QIAquick PCR Purification kit (Qiagen) and were cloned into the pETite C-His vector (Lucigen) with a C-terminal hexahistidine tag. The resulting plasmids were purified using the Monarch Plasmid Miniprep Kit (NEB) and sequenced. The expression plasmids were transformed into *E. coli* Rosetta (DE3) pLysS and were cultured at 37°C shaking overnight in LB broth supplemented with 30 μg mL^-1^ kanamycin (Kan) and 20 μg mL^-1^ chloramphenicol (Cam), then in Terrific Broth (TB) supplemented with the same antibiotics for protein overexpression. Overexpression cultures were grown at 37°C shaking in 1 L of Terrific Broth with Kan and Cam until an OD600 ∼ 0.6 was reached. Isopropyl-β-D-thiogalactopyranoside (IPTG) was added to cultures to induce expression and cells were grown at 30°C shaking for 16- 20 h. Cells were harvested by centrifugation and were stored at −80°C.

### Protein purification

His-tagged BT_1911 and BT_1259 purification from frozen *E. coli* cell pellets was carried out by resuspending pellets in 50 mL of lysis buffer (50 mM NaPO_4_, 300 mM NaCl, 20 mM imidazole, 10 mM 2-mercapoethanol, protease inhibitor (Roche), DNAse (Sigma), pH 8.0). Cells were lysed by sonication. Cell debris was pelleted by centrifugation at 25,000 x *g* for 30 min at 4°C. Lysates were run over a gravity column containing 4 mL of fresh HisPur cobalt resin (Thermo Scientific) equilibrated in wash buffer (50 mM NaPO_4_, 300 mM NaCl, 20 mM imidazole, pH 8.0). Bound BT_1911 was washed on the column with 20 mL of wash buffer and a flow rate of ∼1 mL per min and was eluted with 10 mL of elution buffer, which was wash buffer supplemented with 150 mM imidazole and 10 mM DTT, and flash frozen in liquid N_2_ to prevent oxidation. Enzyme was quantified using the Qubit Protein Assay Kit (Invitrogen) and protein purity was assessed using 4-20% SDS-PAGE gels (Thermo Scientific).

### HSDHa catalytic efficiency assays

BT_1911 kinetics were determined with slight modifications to previously established methods[47]. 0.2 nM HSDH was added to pre-warmed 37°C 100 μL reactions containing 5 mM NAD^+^ in 10 mM MOPS. Bile acid substrates were included in concentrations ranging from 40 μM to 4.5 mM. Change in absorbance at 340 nm was monitored every 80 seconds in flat bottom UV-Star 96 well microplates (Greiner) from a Tecan Infinite F200 Pro plate reader. A standard curve of NADH was included and reaction absorbances were converted to substrate concentration since NADH is generated stoichiometrically with reaction products. Initial velocity data was plotted and Michaelis-Menten parameters were calculated in GraphPad Prism by fitting the data to a nonlinear regression model.

### BSHa activity assay

BSH activity assays were performed as previously described [19]. Briefly, the assay reacted 100 nM BSH with 9 mM bile acids for 60 min. Reactions were carried out in 0.1 mM Na phosphate, 10 mM DTT, pH 6.0 and stopped with 50 μL of trichloroacetic acid. To determine the quantity of amino acid release, a colorimetric ninhydrin reaction was carried out. 25 μL of the quenched BSH reaction was added to 475 μL of ninhydrin buffer (0.3 mL glycerol, 0.175 mL 0.5 M Na Citrate pH 5.5, 0.25% ninhydrin reagent) and boiled for 14 min. A standard curve of the respective conjugated amino acid or 6-APA was prepared for each assay. Absorbance was measured at 570 nm in clear flat-bottom plates in a Tecan Infinite F200 Pro plate reader. No detectable color change was observed indicating that BT_1259.

### Animals and housing

Male and female axenic, C57BL/6J mice (aged 4-9 weeks old) were purchased from the NCSU Gnotobiotic Core (Raleigh, NC) for use in gnotobiotic infection experiments. A biological safety cabinet was cold sterilized with Clidox (CAS Activator: 79-14- 1, Base: 7758-19-2), and used to house all axenic animals and supplies used in experiments. The food, bedding, and water were all autoclaved, and other supplies that were unable to be autoclaved were cold sterilized with Clidox. Female specific pathogen free C57BL/6L mice (aged 4 weeks old) were purchased from Jackson Laboratories for use in antibiotic treated infection experiments. The mice were housed with autoclaved bedding and water, and irradiated food. Cage changes were performed weekly in a laminar floor hood. All mice were subjected to a 12 hr light and 12 hr dark cycle. Animal experiments were conducted in the Laboratory Animal Facilities located on the NCSU CVM campus. The animal facilities are equipped with a full time animal care staff coordinated by the Laboratory Animal Resources (LAR) division at NCSU. The NCSU CVM is accredited by the Association for the Assessment and Accreditation of Laboratory Animal Care International (AAALAC). Trained animal handlers in the facility fed and assessed the status of animals several times per day. Those assessed as moribund were humanely euthanized by CO_2_ asphyxiation. This protocol is approved by NC State’s Institutional Animal Care and Use Committee (IACUC).

### Gnotobiotic mouse colonization experiments

Groups of 9 week old C57BL/6J germ free mice (n=16) were kept inside a sterilized biosafety cabinet for the duration of the experiment. All supplies were sterilized by either cold sterilization by Clidox, or steam sterilization by autoclave. At day 0, all mice except the germ free control group (n=4) were challenged via oral gavage with 10^6^ CFU of *B. theta* WT (n=4), 10^6^ CFU *B. theta* triple KO (n=4), or both 10^6^ CFU *B. theta* WT and 10^6^ CFU *B. theta* triple KO (n=4). All mouse stool tested culture negative for any bacteria before the challenge. Body weight was measured, and fecal pellets were collected throughout the experiment on days 0, 1, 3, and 7. Bacterial enumeration was performed on the fecal pellets plated on BHI-HB. Animals were humanely euthanized by CO_2_ asphyxiation followed by cervical dislocation prior to necropsy. The cecal content was collected at necropsy for bacterial enumeration. Cecal tissue snips were flash frozen in liquid nitrogen and stored at −80C until further analysis.

### Antibiotic treated mouse colonization experiments

Groups of 4 week old female C57BL/6J mice (n=16) were given cefoperazone (0.5mg/mL) in drinking water ad libitum for 5 days. This was followed by a 2 day washout with regular drinking water (Gibco Laboratories, 15230). At day 0, all mice except the control group (n=4) were challenged via oral gavage with 10^6^ cfu of *B. theta* WT, 10^6^ *B. theta* triple KO, or both 10^6^ *B. theta* WT and 10^6^ *B. theta* triple KO. All mouse stool tested culture negative for any bacteria before the challenge. Body weight was measured every day of the experiment, and fecal pellets were collected throughout the experiment on days 0,1,3,5, and 7. Bacterial enumeration was performed on the fecal pellets using Bacteroides Bile Esculin Agar (Anaerobe Systems). Animals were humanely euthanized by CO2 asphyxiation followed by cervical dislocation prior to necropsy. The ileal content, and cecal content were collected at necropsy for bacterial enumeration using Bacteroides Bile Esculin Agar (Anaerobe Systems). Tissue snips were flash frozen in liquid nitrogen and stored at −80C until further analysis.

### Quantification of *B. theta* strains in co-colonization experiments

Genomic DNA was purified from *B. theta* WT and triple KO cultures using Qiagen DNeasy UltraClean Microbial Kit. Genomic DNA from mouse fecal samples was performed using QIAamp® Fast DNA Stool Mini Kit. A standard curve using gDNA from pure culture was generated containing an increasing ratio of *B. theta* WT to triple KO DNA at a known concentration. *bshB* was used to quantify the amount of *B. theta* WT and was normalized to a gene present in both *B. theta* WT and triple KO, 16S in gnotobiotic mice and BT_3075 in antibiotic treated mice. The C_t_ of *bshB* was normalized to the gene present in both bacteria(2^-ΔCt^). This standard curve was then used to quantify the WT to triple KO ratio in mouse fecal, cecal content, and ileal content samples.

### Statistical analysis

With the exception of the RNAseq analysis, all statistical tests were performed in GraphPad Prism 8. Ordinary two-way ANOVA with Tukey’s multiple comparisons test was used to determine statistical differences between conjugated and deconjugated bile acids, while Mann-Whitney was used to determine differences between strains in the MIC datasets. Ordinary two-way ANOVA with Dunnet’s multiple comparison test was used to determine conditions statistically different from the no bile acid control, while ordinary two way ANOVA with Šídák’s multiple comparisons test was used to determine differences between strains in PI datasets. Ordinary one-way ANOVA with Dunnett’s multiple comparisons test was used on log transformed data to determine conditions significantly different from the parental strain in CFU growth experiments. Unpaired t test with Welch’s correction was used to determine conditions significant differences between mid-log and stationary phase, two-way ANOVA with Tukey’s multiple comparison test was used to determine differences in expression of genes at mid log in qPCR data. The R package DESeq2 used Wald tests with Benjamini & Hochberg correction between strains and Bonferroni correction between bile acids to identify statistically significant differentially expressed genes in the RNASeq datasets. Ordinary two-way ANOVA with Tukey’s multiple comparison test were used to determine significant differences between log transformed CFU/g of different strains in the mouse model. One sample t-test was used to determine significant if WT strain was significantly different from a theoretical mean of 50% within the competition mouse model. Statistical significance is displayed as follows, * p<0.05; ** p<0.01; *** p<0.001; **** p<0.0001.

## Data availability

Raw sequences have been deposited in the Sequence Read Archive (SRA) under BioProject ID: PRJNA986925. Source data are provided within each supplementary data file. Other data and biological materials are available from the corresponding author upon reasonable requests.

## Results

### *B. theta* is more sensitive to deconjugated bile acids that alter membrane integrity

To address if and how *B. theta*’s bile acid altering enzymes are beneficial to the bacterium, seven mutant strains were generated by knocking out the genes encoding bile acid altering enzymes *bshA* (BT_1259), *hsdhA* (BT_1911), and *bshB* (BT_2086) individually and in combination. Gram-negative bacteria are thought to be inherently more resistant to bile acids than Gram-positive bacteria as they are frequently used in media to select for Gram-negative bacteria [48]. Therefore, we hypothesized that *B. theta* would be able to grow in high concentrations of bile acids. To address this, we grew *B. theta* WT in increasing concentrations of TCA, GCA, CA, TCDCA, GCDCA, CDCA, TDCA, GDCA, and DCA to determine its minimum inhibitory concentration (MIC). Since WT *B. theta* has the ability to modify bile acids (Figure 1A), a strain of *B. theta* with all three genes encoding bile acid altering enzymes knocked out (*ΔbshA ΔhsdhA ΔbshB* referred to in this paper as the *B. theta* triple KO) was also tested against the same panel of bile acids. *B. theta* was inhibited by higher concentrations of TCA (16.6 mM), GCA (16.6 mM), and CA (10 mM), while *B. theta* was more sensitive to TCDCA (2.5 mM), GCDCA (2.5 mM), CDCA (0.625 mM), TDCA (1.66 mM), GDCA (1.25 mM), and DCA (0.625 mM) (Figure 1B). Overall, *B. theta* was only able to grow in approximately half the amount of deconjugated bile acids (CA, CDCA, and DCA) when compared to their conjugated forms (T/GCA, T/GCDCA, T/GDCA) (Figure 1B). The differences between conjugated and deconjugated were significant between T/GCA and CA in both strains (p<0.0001 by two-way ANOVA with Tukey’s multiple comparison, Suppl Figure 1A). The *B. theta* triple KO strain behaved similarly, with the only significant difference being between conjugated and deconjugated forms of T/GCA and CA (p<0.0001 by two-way ANOVA with Tukey’s multiple comparison, Suppl Figure 1B). The only difference in MICs between the WT and the triple KO strain was in the presence of GDCA (2.5 mM vs 1.25 mM, p<0.05 by Mann-Whitney, Figure 1B).

**Figure 1.**
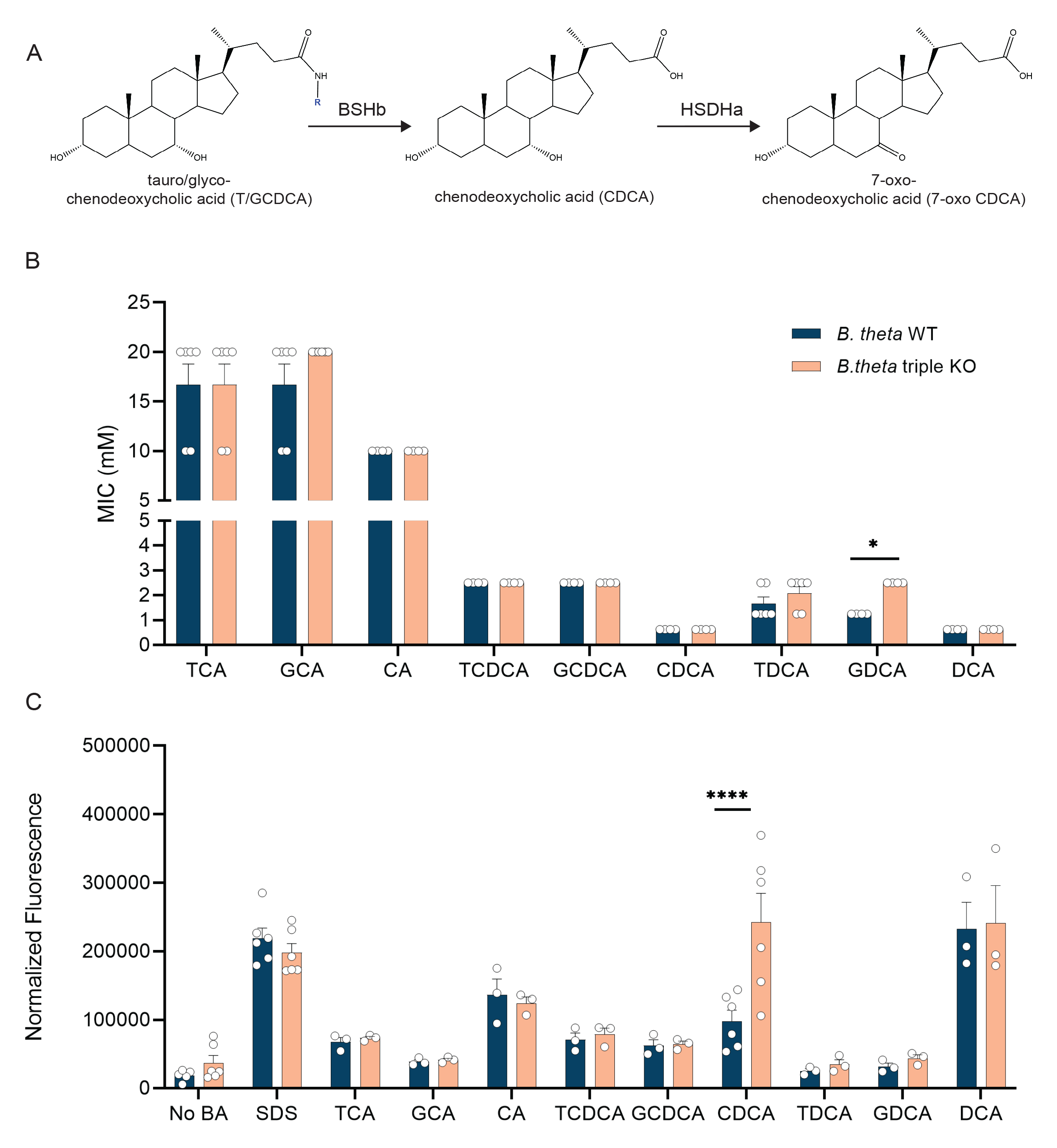
*B. theta* is more sensitive to deconjugated bile acids which are able to alter the membrane integrity. (A) Example of bile acid transformation carried out by BSHb and HSDHa. (B) WT and the triple KO strains of *B. theta* were used to determine MICs in different bile acids. Bars represent mean MICs from (n = 2-3 biological replicates, n=2 technical replicates). Error bars represent standard error. Asterisks denote significant (p<0.05) by Mann-Whitney between strains. (C) WT and the triple KO strains were incubated with different bile acids and stained with propidium iodide. Asterisks denote significant (p<0.0001) differences between strains by 2-way ANOVA with Šídák’s multiple comparisons test.

Using propidium iodide staining we next evaluated how each bile acid affected the membrane integrity of both the WT and triple KO strain. When compared to a no bile acid control only deconjugated bile acids, including CA, CDCA, and DCA, were able to significantly alter the membrane in both *B. theta* WT and *B. theta* triple KO (Suppl Figure 1C and D, p<0.05, p<0.01, p<0.0001 by two-way ANOVA with Dunnet’s multiple comparisons test). The only major difference between strains was in the presence of CDCA, where the triple KO was significantly more sensitive to membrane damage compared to the WT (p<0.0001 by two-way ANOVA with Šídák’s multiple comparisons test, Figure 1C). To determine if one gene is responsible for this phenotype, strains of *B. theta* with single gene deletions with CDCA in a membrane integrity assay. *B. theta* triple KO had significantly higher membrane damage than *B. theta ΔbshA,* and *B. theta ΔbshB,* but was not higher than *B. theta ΔhsdhA* (p<0.05, p<0.001 by two-way ANOVA with Šídák’s multiple comparisons test, Suppl Figure 1E), suggesting the missing *hsdhA* is contributing to the phenotype.

### *bshB* decreases survival in the presence of conjugated forms of CDCA and DCA

Given the differences in both the MIC and membrane integrity assays between the *B. theta* WT and triple KO strains we wanted to better understand how each gene encoding the bile acid altering enzymes affects growth alone and in combination with the others. A subinhibitory concentration of each bile acid was selected based on their individual MIC. Since some bile acids increased lag phase growth of the WT strain (Suppl Figure 2), we evaluated bacterial growth via enumeration at 12 and 24 hours to capture differences in growth kinetics (Figure 2 and Suppl Figure 3).

**Figure 2.**
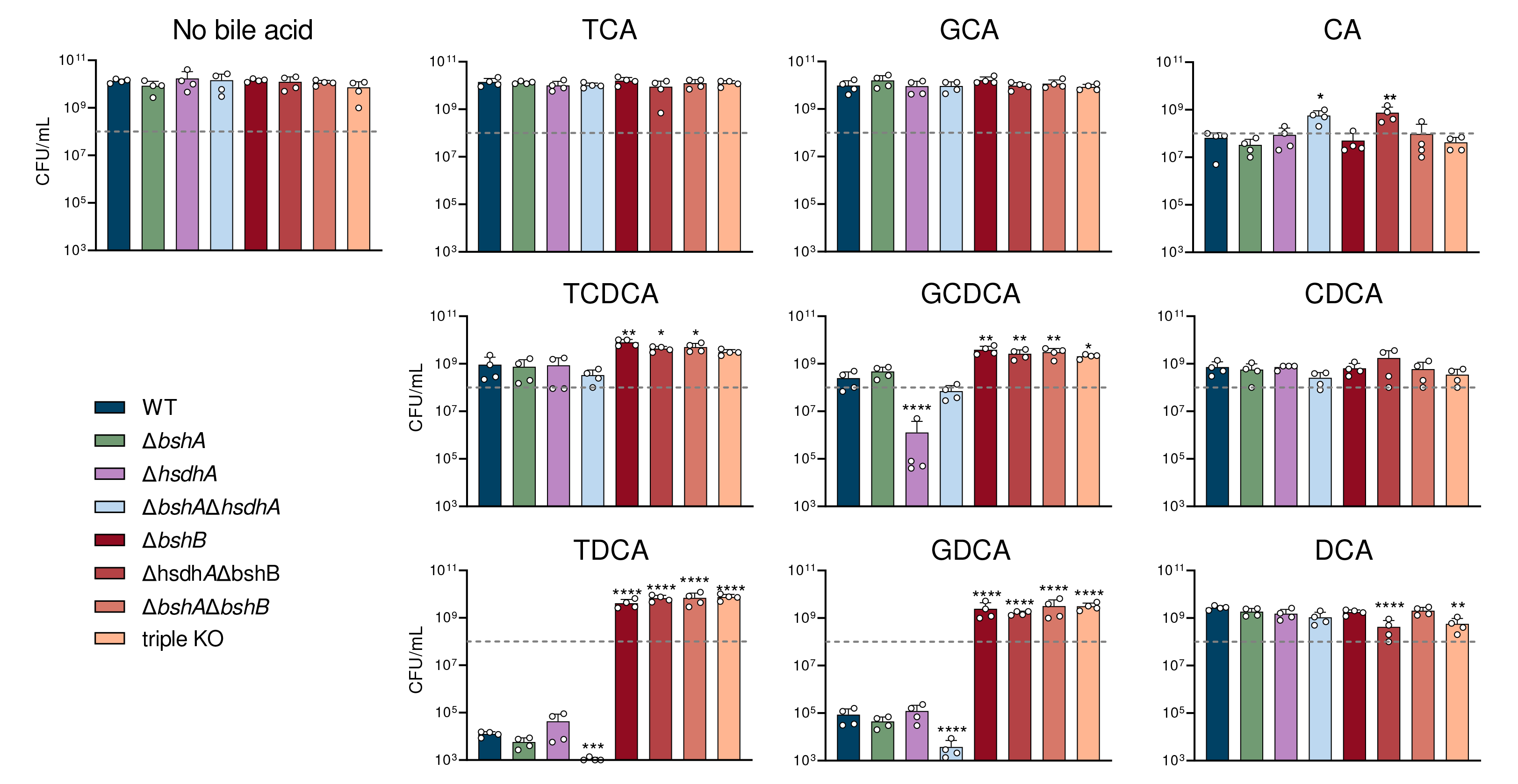
Genes encoding bile acid altering enzymes have differential effects on *B. theta*’s fitness. Bacterial enumeration of *B. theta* WT, Δ*bshA*, Δ*hsdhA,* Δ*bshA*Δ*hsdh,* Δ*bshB,* Δ*hsdhA*Δ*bshB*, Δ*bshA*Δ*bshB*, and triple KO strains after 24 hr of growth in TYG supplemented with different bile acids. Error bars represent standard deviation from n≥2 biological replicates. Dashed grey line represents the approximate starting CFU/mL added at 0 hr. Asterisks denote significant (* p<0.05; ** p<0.005; *** p<0.0005; **** p<0.00005) differences from WT strain by one way ANOVA.

In the no bile acid control, all strains grew to similar levels after 24 hours of growth (Figure 2). In accordance with the lack of BSHa activity seen previously [23], Δ*bshA* had a similar growth profile to the WT strain in all bile acid conditions (Figure 2). We also heterologously expressed and purified BSHa from *E. coli* (Suppl Figure 4, BT_1911 is HSDHa and BT_1259 is BSHa), and it did not show any BSH or penicillin V acylase activity *in vitro*. However, strains lacking a *bshB* were significantly more resistant to TCDCA, GCDCA, TDCA and GDCA toxicity relative to the WT strain (p<0.05, p<0.01, p<0.001, p<0.0001 by one way ANOVA with Dunnet’s multiple comparison test, Figure 2). Strains containing *bshB* had growth that trended lower than strain’s missing *bshB* (Figure 2), further supporting the role of BSHb as the only active BSH encoded by *B. theta*. These findings suggest that *bshB* may be detrimental to *B. theta’s* growth in the presence of TCDCA, GCDCA, TDCA and GDCA.

### *B. theta* hydroxysteroid dehydrogenase (HSDHa) acts on CDCA core bile acids

While HSDHs play an important role in the metabolism of secondary bile acids [49], they have not been directly implicated in bacterial fitness. However, Δ*hsdhA* survival was significantly reduced when grown in the presence of GCDCA when compared to the WT strain (p<0.0001 by one way ANOVA with Dunnet’s multiple comparison test, Figure 2). Similar increases in survival were observed in the absence of the BSHa and BSHb individually as well (Figure 2), suggesting that HSDHa may be acting on GCDCA in addition to its deconjugated form, CDCA. Since there are few examples of conjugated bile acid recognition by HSDHs in the literature [35, 50, 51], we characterized the activity and substrate range of HSDHa by heterologously expressing and purifying the enzyme to homogeneity from *E. coli* (Suppl Figure 4). HSDHa displayed high activity on the bile acids CA, CDCA, GCDCA, and TCDCA measured via the reduction of cofactor NAD^+^ to NADH using a UV-visible spectrophotometer, however it displayed a clear preference for CDCA (Table 1). Furthermore, the catalytic efficiency of GCDCA dehydrogenation remained high despite the glycine conjugation (Table 1). These *in vitro* observations support a mechanism of HSDHa dehydrogenation of GCDCA during *B. theta* growth. While it is not clear if this transformation reduces bile acid toxicity or adapts metabolism for persistence, taken together these data suggest that *hsdhA* promotes survival in the presence of GCDCA.

**Table 1.**
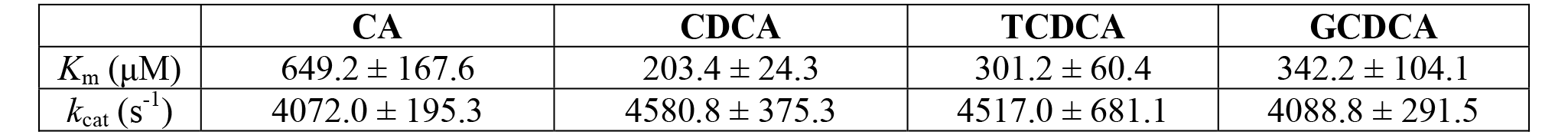
Catalytic efficiency measurements of the 7-α HSDH encoded by *B. theta*.

|  | CA | CDCA | TCDCA | GCDCA |
| --- | --- | --- | --- | --- |
| $K_m$ (μM) | 649.2 ± 167.6 | 203.4 ± 24.3 | 301.2 ± 60.4 | 342.2 ± 104.1 |
| $k_{cat}$ ($s^{-1}$) | 4072.0 ± 195.3 | 4580.8 ± 375.3 | 4517.0 ± 681.1 | 4088.8 ± 291.5 |

### Bile acid altering enzymes are highly expressed in stationary phase of growth when nutrients are limited

To gain further insight into how bile acid altering enzymes impact *B. theta*’s growth in subinhibitory concentrations of bile acids, their expression was monitored by qRT-PCR. RNA was extracted at mid-log and stationary phase from rich medium (TYG) cultures supplemented with each bile acid. Expression of *bshA* increased in stationary phase of growth compared to mid-log phase of growth in all bile acids except for CA and DCA (p<0.05, p<0.01, p<0.001 by unpaired t-test with Welch’s correction, Figure 3A). There were no significant differences in the expression of *bshB* between mid-log and stationary phase growth (Figure 3B). The variability between the replicates could be due to an active BSHb, but a similar trend can still be observed, where expression increases in stationary phase (Figure 3B). Expression of *hsdhA* increased significantly in stationary phase in the presence of CDCA, TDCA, and GDCA (p<0.05, p<0.01 by unpaired t-test with Welch’s correction) while decreasing in the presence of CA (p<0.01 by unpaired t-test with Welch’s correction, Figure 3C). Additionally, expression of *hsdhA* was higher than *bshA* and *bshB* in both CA and GCDCA during mid-log growth (p<0.00005 by two-way ANOVA with Tukey’s multiple comparison test, Suppl Figure 5). This high level of expression during stationary phase may be reflective of the nutrient deprived or stressed environment.

**Figure 3.**
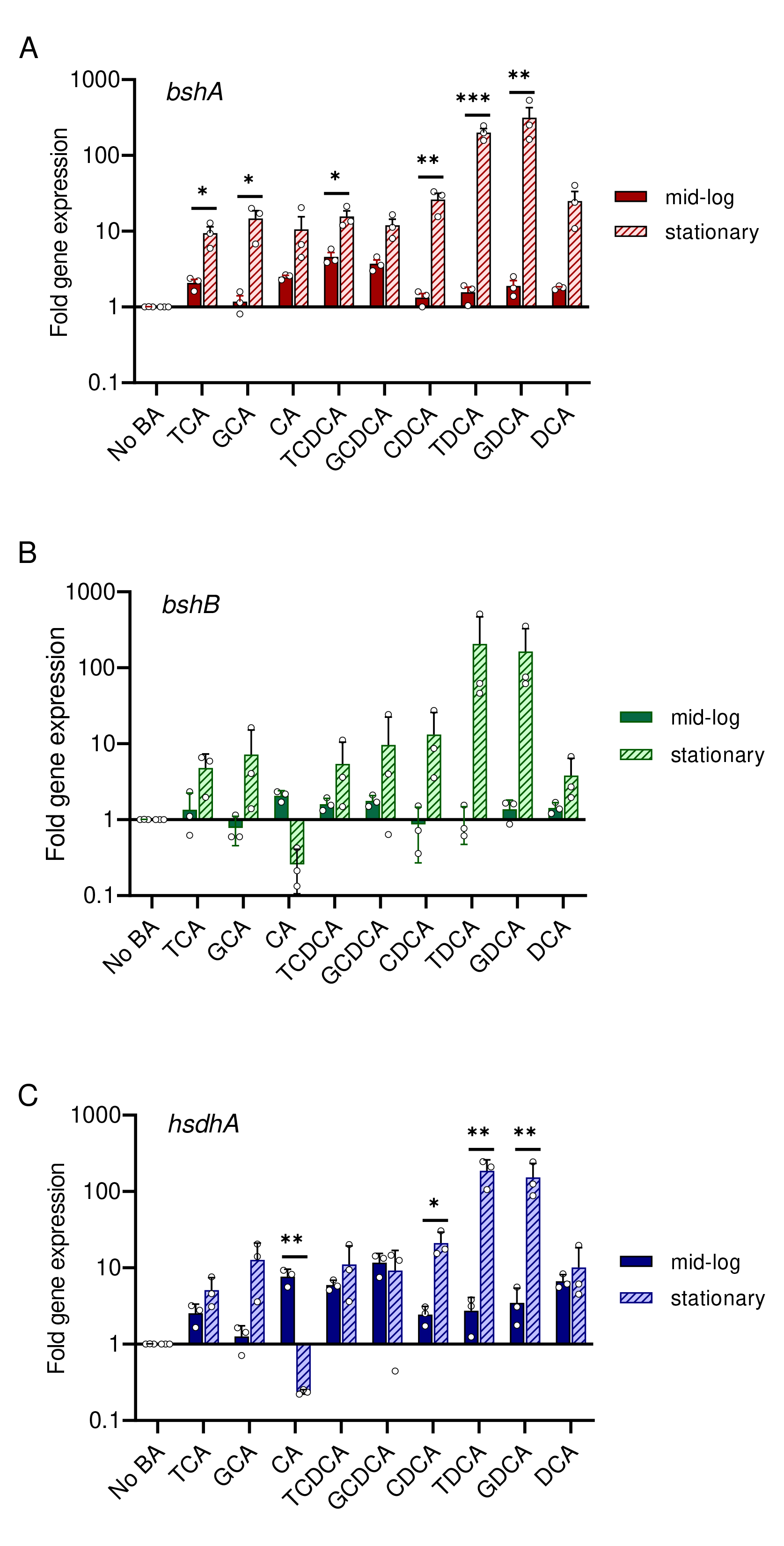
Genes encoding bile acid altering enzymes are highly expressed in stationary phase of growth when nutrients are limited. Bars represent fold change in gene expression for (A) *bshA,* (B) *bshB*, (C) *hsdhA* from mid-log and stationary phase cultures grown in different bile acids compared to growth in the absence of bile acid. Data was normalized using the ΔΔCt method using 16S rRNA as a housekeeping gene. Error bars represent standard error between replicates (n=3 biological, 3 technical replicates each). Asterisks denote significant (* p<0.05; ** p<0.01; *** p<0.001) differences between mid-log and stationary phase growth stages by t-test.

### Metabolism of *B. theta* is impacted by bile acid altering enzymes

Given that genes encoding bile acid altering enzymes are highly expressed in stationary phase of growth when nutrients are limited, we sought to determine the effect that the bile acid altering enzymes have on the global metabolic transcriptome in nutrient limiting conditions. RNA was extracted and purified from WT and triple KO cultures grown to mid-log phase growth in rich (TYG) and minimal media, which has reduced glucose and no tryptone or yeast extract, and differential expression analysis was performed. Differential expression analysis was performed to determine how each strain responds to nutrient limiting conditions, then we compare these responses between the strains. The total number of genes that are increased in nutrient limited conditions due to the presence of bile acid altering enzymes was 117 (Up), while the number of genes that decrease in response to nutrient limited conditions due to the presence of bile acid altering enzymes was 40 (Down) (Figure 4A). These are genes that changed in expression in nutrient limitation due to the genotype. To gain insight into the types of genes differentially expressed, we determined KEGG orthologs. We were able to successfully identify KEGG orthologs for 10 genes positively associated with bile acid altering enzymes in nutrient limited conditions and 17 genes negatively associated with bile acid altering enzymes in nutrient limited conditions (Figure 4B). Carbohydrate metabolism and amino acid metabolism were the most affected metabolic pathways (Figure 4B). Lipid and energy metabolism were also impacted to a lesser degree. The genes identified in carbohydrate metabolism are involved in a variety of pathways. The response to nutrient limitation due to the presence of bile acid altering enzymes shows a reduction in expression of genes involved in the metabolism of specific sugars such as galactose, glucosamine, and glutamine, whereas there is an increase in broader enzymes including biotin carboxylase, pyruvate dehydrogenase, and alpha-glucosidase (Figure 4C). Amino acid metabolism was impacted primarily through an operon BT_3757-BT_3760, which is involved in arginine/ornithine metabolism, and was reduced in expression in nutrient limited conditions due to the presence of bile acid altering enzymes (Figure 4C). A serine acetyltransferase and L-asparaginase increased in response to nutrient limitation due to the presence of bile acid altering enzymes was also observed (Figure 4C). Many genes within lipid metabolism are also categorized as carbohydrate metabolism with the only unique gene being a long chain fatty acid CoA ligase which decreased in expression due to the presence of bile acid altering enzymes (Figure 4C). These genes may interact with bile acids as bile acids help dissolve lipids during digestion. Energy metabolism, which HSDHs may play a role in the electron transport chain, show a decrease in cytochrome C552 precursor and an increase in oxidoreductase, 2-nitropropane dioxygenase family due to the presence of bile acid altering enzymes (Figure 4C).

**Figure 4.**
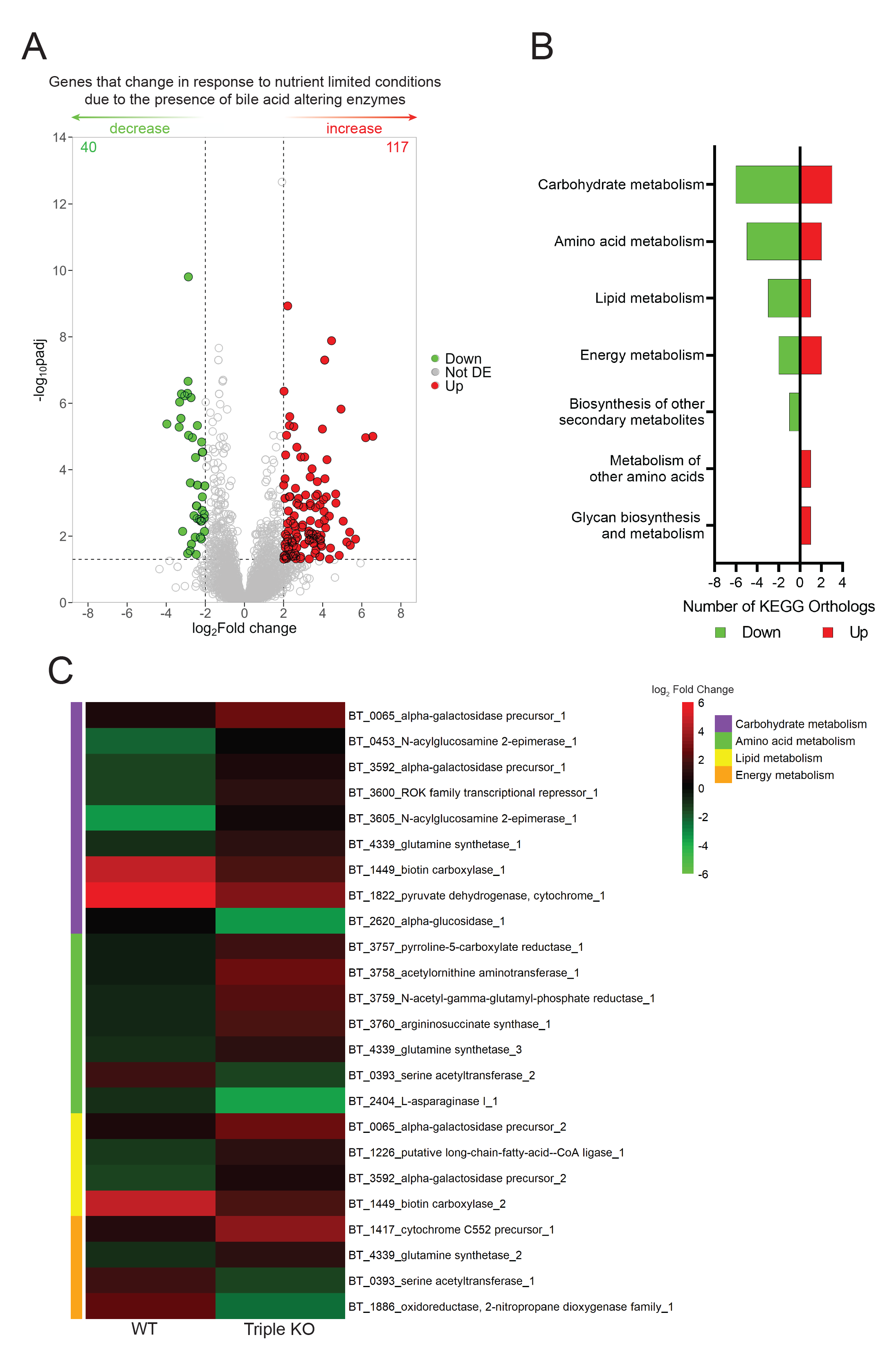
Global metabolism of *B. theta* is impacted by the absence of bile acid altering enzymes in nutrient limited conditions. (A) Volcano plot of differentially expressed genes identified by RNASeq during nutrient limiting conditions. Points are colored when p < 0.05 by Wald test with Benjamini and Hochberg correction and log_2_fold change is > 2 (red) or < 2 (green). The number of significant genes is listed in the top corners for both directions. (B) KEGG orthologs detected for genes listed in (A). (C) Heatmap of differentially expressed genes present in the top four KEGG orthology categories in (B).

### Carbohydrate metabolism is most affected by bile acid exposure in nutrient limited conditions

Since carbohydrate and amino acid metabolism were most affected in Figure 4, and amino acids are liberated by BSHs, we sought to determine how bile acids affect the global metabolic transcriptome of WT *B. theta* in nutrient limited conditions. RNA was purified from WT cultures grown in rich or minimal media in the presence and absence of the following bile acids; TCA, GCA, CA, TCDCA, GCDCA, CDCA, TDCA, GDCA, DCA. Differential expression analysis was done to determine which genes respond to nutrient limiting conditions in the presence of bile acids, compared to a no bile acid control. The number of genes differentially expressed varied by bile acid, with DCA having the most differentially expressed genes, with 59 increased in expression and 13 decreased in expression (Suppl Figure 6). A large response to polysaccharide utilization locus (PUL) associated genes and carbohydrate metabolism was observed in the presence of most bile acids (Figure 5A, B, Suppl Figure 6). The majority of genes were increased in expression in nutrient limited conditions due to the presence of a bile acid. A similar trend was observed in genes associated with the metabolism of cofactors and vitamins (Figure 5C, Suppl Figure 7), with the exception of TCDCA and GCDCA. Interestingly, the addition of a bile acid decreased membrane transport in nutrient limiting conditions (Figure 5D, Suppl Figure 7), potentially implicating some genes as bile acid transport systems. Signal transduction increased in nutrient limiting conditions due to the presence of bile acids (Figure 5E, Suppl Figure 7). Amino acid metabolism was minimally affected by the presence of bile acids, and no correlation between amino acid metabolism and bile acids for which *B. theta* BSHb has activity on, was observed (Figure 5F, Suppl Figure 7).

**Figure 5.**
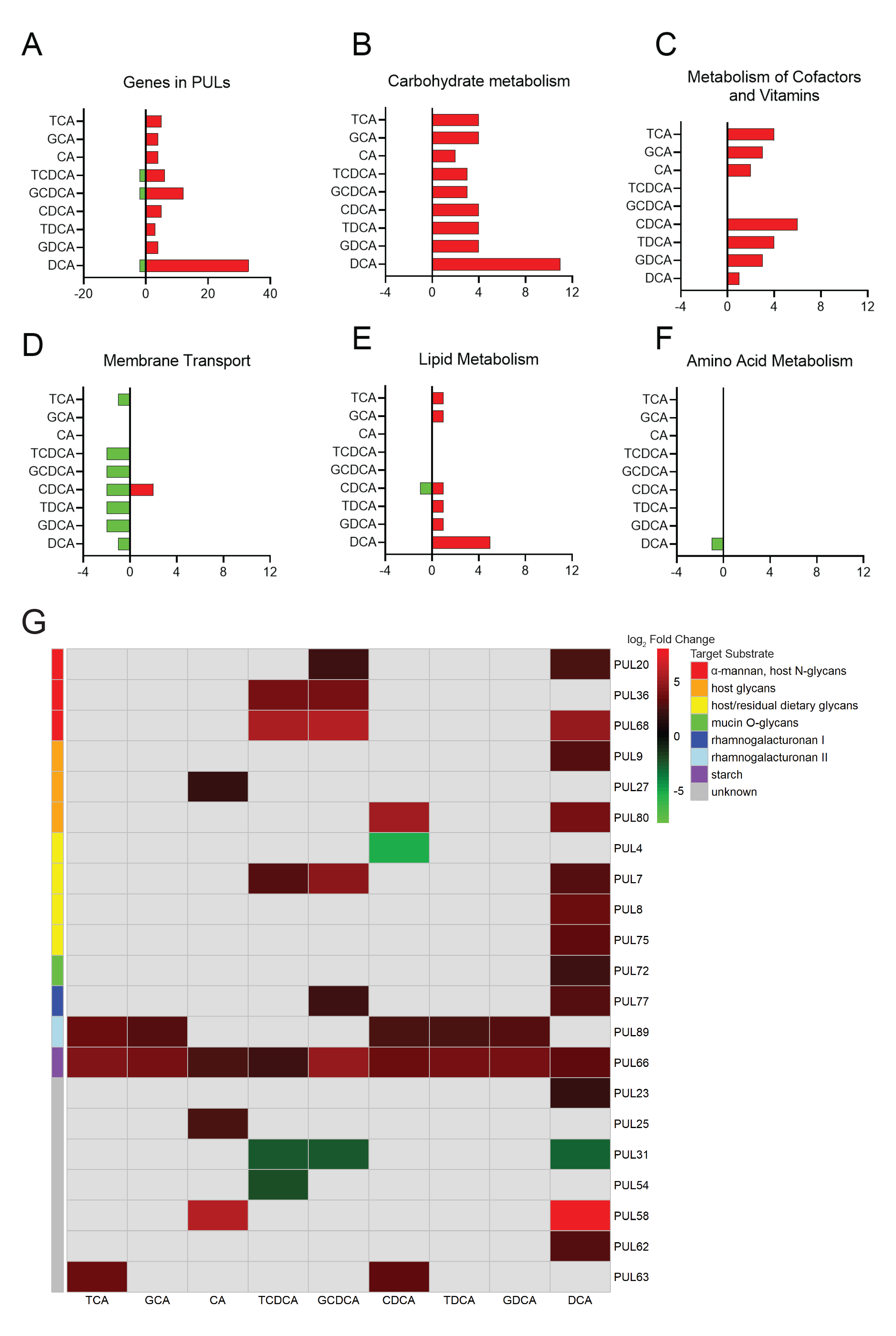
Different bile acids are able to alter *B. theta*’s metabolism in nutrient limiting conditions. Differently expressed genes that fall under KEGG orthologs: (A) PUL and (B) carbohydrate metabolism, (C) metabolism of cofactors and vitamins, (D) membrane transport, (E) lipid metabolism, and (F) amino acid metabolism. Genes are displayed that had expression of p < 0.05 by Wald test with Bonferroni correction and log_2_fold change is greater than 2 or less than −2. (G) Heatmap of PUL genes that were differentially expressed in nutrient limited conditions due to the presence of a bile acid. Target substrates and gene annotation of PULs were annotated using CAZy’s PULDB (http://www.cazy.org/PULDB/) and from multiple publications for deeper annotation [39-46].

*B. theta*’s response to nutrient limitation was impacted differently depending on the sterol core and amino acid conjugate, in both PULs (Figure 5G) and other types of metabolism (Suppl Figure 7). Of the 21 PULs that had genes differentially expressed, DCA showed the broadest ability to impact them by affecting 14 PULs (Figure 5G). Bile acids primarily increased expression of genes in PULs in nutrient limited conditions with the exceptions being PUL4, PUL31, and PUL54 that decreased in expression due to the presence of some bile acids (Figure 5G). PUL66, implicated in *B. theta*’s ability to degrade starch, was most broadly impacted by all bile acids, which was independent of sterol core or conjugated amino acid (Figure 5G). All genes within each PUL were not differentially expressed consistently, with some genes within the PUL responding more than others (Suppl Figure 8). Neither taurine nor glycine metabolism genes were impacted by the addition of bile acids (Suppl Figure 9), but this could be due to the limited annotation in KEGG as well.

### Bile acid altering enzymes in *B. theta* are able to impact colonization in a mouse model

To further understand the benefit that *B. theta*’s bile acid altering enzymes confer we wanted to examine their contribution to bacterial fitness in the presence of a complex bile acid pool in a host or mouse model. Germ-free mice were monocolonized with either WT, the triple KO, or co-colonized with both strains to define colonization and competition over a 7-day period (Figure 6A). Mice monocolonized with *B. theta* WT and co-colonized with both strains had a significantly higher fecal bacterial load compared to mice monocolonized with *B. theta* triple KO at day 1 post challenge, (p<0.05, p<0.001 by two-way ANOVA with Tukey’s multiple comparison, Figure 6A). There were no differences in bacterial load in the feces at day 3 or 7 or in day 7 cecal content (Figure 6A). However, when co-colonized, the WT strain had a significant competitive advantage compared to the triple KO at day 7 in fecal samples (p<0.05 by one sample t-test, Figure 6B).

**Figure 6.**
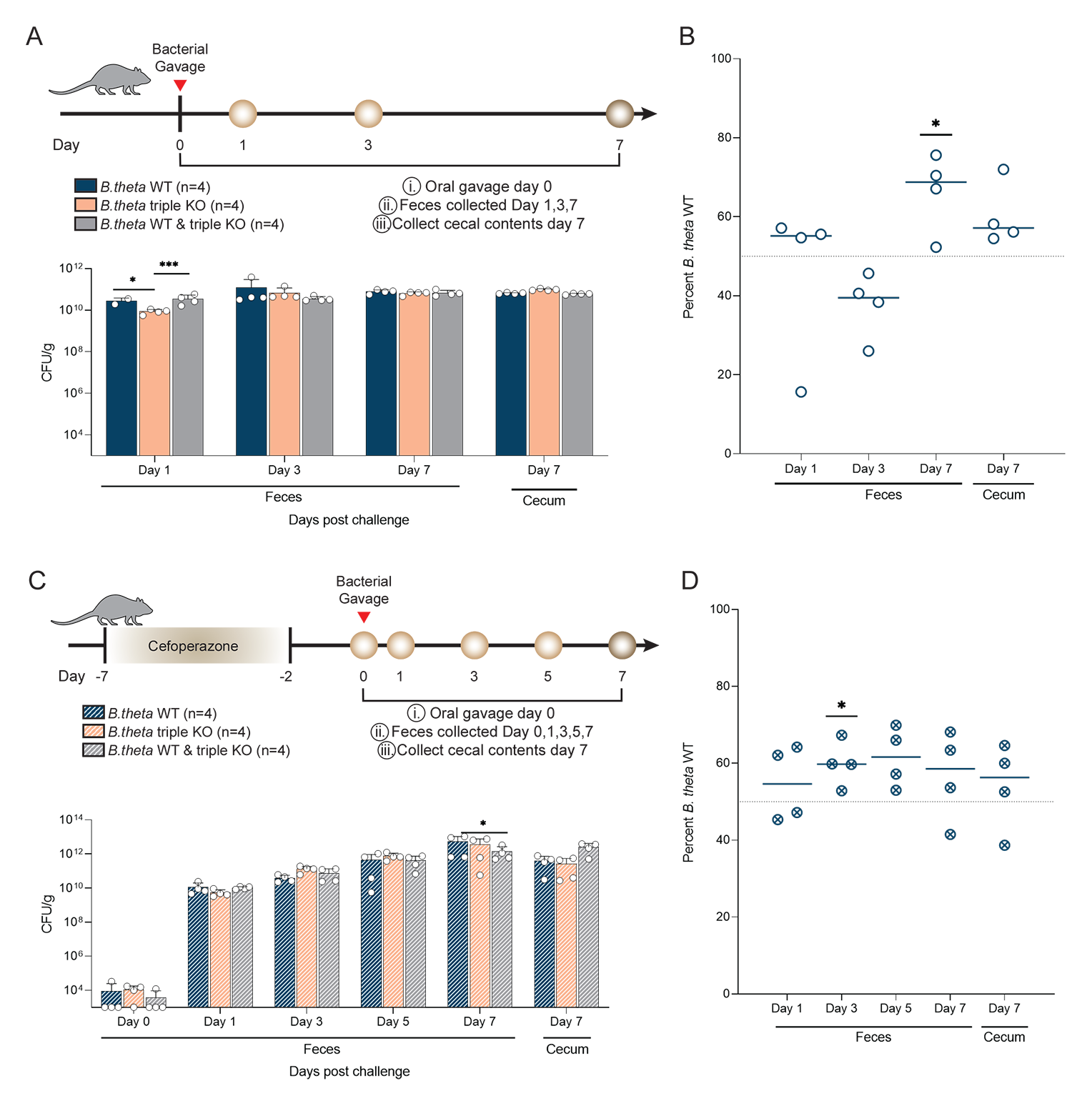
Genes encoding bile acid altering enzymes impact colonization in a mouse model. (A) Experimental design for gnotobiotic mouse model. Bacterial enumeration of the feces and cecal content in mice monocolonized with *B. theta* WT (n=4), triple KO (n=4), or mice co-colonized with an equal amount of *B. theta* WT and triple KO (n=4). Asterisks denote significant (p<0.05, p<0.001) by two-way ANOVA with Tukey’s multiple comparison test. (B) qPCR of DNA extracted from fecal and cecal content of gnotobiotic mice. Asterisks denote significance (p<0.05).via one sample t test with a theoretical mean of 50%. (C) Experimental design for antibiotic treated mouse model. Bacterial enumeration of the feces and cecal content of cefoperazone treated mice inoculated with *B. theta* WT (n=4), triple KO (n=4), or mice co-inoculated with an equal amount of *B. theta* WT and triple KO (n=4). Asterisks denote significance (p<0.05) by two-way ANOVA with Tukey’s multiple comparison test. (D) qPCR of DNA extracted from fecal or cecal content. Asterisks denote significant (p<0.05) one sample t test with a theoretical mean of 50%.

In order to measure the bacterial fitness of these strains in a more complex host, where the microbiota is intact and the bile acid pool is even more complex, we leveraged an antibiotic treated mouse model. Cefoperazone treated mice were challenged with either WT, the triple KO, or both strains at Day 0 (Figure 6C). Antibiotic treated mice challenged with both strains had higher bacterial load in the cecum at day 7 compared WT and triple KO strain alone (p<0.05 by two-way ANOVA with Tukey’s multiple comparison, Figure 6C). Similarly, the WT strain had a significant growth advantage on day 3 in fecal samples, but this advantage was lost in later timepoints potentially due to the returning microbiota (p<0.05 by one sample t-test Figure 6D).

## Discussion

Bile acids are host synthesized molecules that are modified by the gut microbiome. These modifications impact the host via immunity, metabolism, and the circadian rhythm [23] while also having an impact on the microbiota [19, 52]. Increasing our understanding of how members of the gut microbiota are able to perform these biotransformations on bile acids and how this benefits the bacterium and surrounding community will help aid in efforts to leverage the bile acid pool to treat many diseases. *B. theta*, a predominant member of the gut microbiota, encodes three enzymes suspected to modify bile acids; BSHa, BSHb, and HSDHa [28]. Previous work has focused on how these enzymes impact host physiology, but little has focused on how they benefit the bacterium itself. This study hypothesized that *B. theta* uses its BSHs and HSDH to modify bile acids to provide a fitness advantage for itself *in vitro* and *in vivo*.

Historically microbiologists have used bile acids or oxgall in media to select for Gram-negative members of the gut microbiota, so we thought that Gram-negative bacteria would be more resistant to bile acids [48]. When compared to Gram-positive microbes *Lactobacillus gasseri* and *Lactobacillus acidophilus* in a previous study*, B. theta* was much more sensitive to bile acids TCDCA, TDCA, and DCA while being similarly resistant to TCA and GCA [19]. In addition lactobacilli did not show the same pattern of membrane damage, where deconjugated bile acids impacted membrane integrity more than conjugated bile acids [19]. *B. theta* WT had significantly less membrane damage than *B. theta* triple KO in the presence of CDCA. This could indicate that either the HSDHa is mitigating this damage by converting CDCA to 7-oxo lithocholic acid, or the BSHb is converting CDCA to a conjugated form of CDCA ([14-16, 33]).

Prior studies suggested that bacteria that encode BSHs are able to colonize the harsh gut environment better as BSHs deconjugate more toxic conjugated bile acids, making less toxic deconjugated bile acids. Our results do not support this notion and suggest that interactions between gut bacteria that encode different bile altering enzymes and bile acids is more complex and context dependent then we first thought [19]. Deconjugated bile acids, moreover than conjugated bile acids, were able to alter *B. theta*’s growth and this was associated with impaired membrane integrity.

We went on to further characterize *B. theta*’s growth at 12 and 24 hours leveraging a panel of single, double, and triple KO mutants and found that *bshB* was detrimental to bacterial growth in the presence of GCDCA, TDCA and GDCA. BSHb is the primary enzyme associated with deconjugation of conjugated bile acids in *B. theta* and has activity on conjugated forms of DCA, is minimally active on conjugated forms of CDCA, and inactive on conjugated forms of CA [23]. However, this activity was observed in whole cell cultures over a 48 hour period. Purified BSHb has shown limited specific activity over a short period of time [14]. A recent study identifying amino acid residues required for BSH substrate selectivity found that *B. theta* BSHb is missing the selectivity loop that allows for taurine and glycine-specificity [14]. This may reflect a broader ability to deconjugate bile acids that are both taurine and glycine conjugated. The presence of BSHb in this study had the greatest impact on *B. theta*’s growth in the presence of bile acids that it has the highest activity on. The absence of this enzyme did provide a fitness advantage in the presence of GCDCA, TDCA, and GDCA. This could reflect the fact that DCA is more toxic than TDCA and GDCA in Figure 1B, as well as BSHb acts on both TDCA and GDCA [23]. TDCA and GDCA may be being converted into the more toxic form of DCA by this enzyme *in vitro*. Deconjugation of bile acids are detrimental to *B. theta* and this may explain why BSHb is secreted away from the cell in extracellular vesicles [32]. The extracellular secretion of BSHb may also explain why *B. theta* does not utilize taurine or glycine liberated from the bile acids it deconjugates (Suppl Figure 9). We also observed that CDCA and DCA decreased growth in many strains lacking *bshB* (Figure 2)*. B. theta* was able to reconjugate DCA with a variety of amino acids in whole cell cultures [33]. This suggests that BSHb could be reconjugating DCA with different amino acids, into a less toxic conjugated form that protects *B. theta* from further damage.

The loss of *bshA* showed a distinctly different response. Previous work has been unable to define the activity of BSHa in 50 μM of bile acid [23], however knockouts of BSHb show some ability to deconjugate TCA and GCA at higher bile acid concentrations of 5 mM [32]. In addition, this enzyme does not have a selectivity loop that is specific for taurine and glycine conjugated bile acids. We purified BSHa and tested its ability to deconjugate TCA, GCA, TCDCA, GCDCA, TDCA, and GDCA, but were unable to measure activity (data not shown). There were no notable changes to *B. theta*’s tolerance to bile acids, and growth in bile acids due to the absence of this enzyme.

Over the course of the growth experiments *hsdhA* was able to provide a protective effect in presence of GCDCA. Since this enzyme has not be fully characterized we wanted further investigate its substrate specificity and activity in the presence of TCA, GCA, CA, TCDCA, GCDCA, and CDCA. The 7α-hydroxysteroid dehydrogenase encoded by *B. theta* has been previously described to have activity on taurine and glycine conjugated, as well as deconjugated forms of CA and CDCA using concentrated lysates ([35]). *B. theta*’s HSDHa had appreciable activity on CA, as well as taurine or glycine conjugated and deconjugated forms of CDCA (Table 1). The ability to act on conjugated forms of bile acids contradicts the notion that bile acids are first deconjugated before modifications to the hydroxyl group occurs [53, 54]. We also saw that of the three genes encoding bile acid altering enzymes, *hsdhA* is more highly expressed in the presence of CA, TCDCA, GCDCA, and DCA (Suppl Figure 4). HSDHa may be providing a protective effect by changing the structure of the conjugated bile acid before BSHb is able to deconjugate it, where that deconjugation is detrimental to *B. theta’s* growth.

The absence of bile acid altering enzymes led to global metabolic changes in nutrient limited conditions independent of bile acid supplementation. We saw large shifts in response to nutrient limited conditions including genes important for carbohydrate metabolism, amino acid metabolism, lipid metabolism, and energy metabolism between the WT and triple KO strain. Since nutrient limiting conditions lacked amino acids, we expected changes in amino acid metabolism. We did observe a decrease in arginine/ornithine metabolism in nutrient limited conditions due to the presence of bile acid altering enzymes. Ornithine metabolism is important for gut pathogen colonization including *C. difficile* physiology and pathogenesis [55, 56]. This suggests that encoding bile acid altering enzymes could have far reaching metabolic impacts that not only affect the bacterium that encodes them, but gut pathogens as well. The addition of bile acids greatly impacted *B. theta*’s global metabolic response to nutrient limitation and this was dependent on the sterol core and conjugated amino acid. Many carbohydrate utilization genes were increased in expression in nutrient limited conditions due to the presence of a bile acid. Bile acid stress has been shown *in vivo* to alter the microbial metabolism, including carbohydrate metabolism, in the cecum [57]. However, the impact bile acids have on each member of the microbiota has not been well studied.

We also investigated the potential for the amino acid being freed by *B. theta* BSHs, taurine or glycine, being used as a nutrient, or for the electron lost during dehydroxylation to feed into energy metabolism. However, we primarily observed an increase in genes important for carbohydrate metabolism and associated with PULs, not amino acid or energy metabolism. This could be due to the limited annotated genes in that pathway. The primary carbohydrate source used in both the minimal and rich media is glucose. Bile acids directly interact with carbohydrates as they can bind to fiber, with certain fiber sources such as lignin binding more so than bran [58]. Fiber intake also has impacts on the composition of the bile acid pool [59]. Considering the strong response in carbohydrate metabolism in the presence of bile acids, *B. theta* may be using different carbohydrates to exploit bile acid-fiber binding and, thus altering the types of fiber available in the gut. If bile acids are bound to fiber, they can’t be reabsorbed through enterohepatic circulation and the host’s homeostasis will synthesize more bile acids from cholesterol [60]. This has been proposed as a way to reduce cholesterol, and our findings may indicate that the host microbiome, more specifically *B. theta* might influence this interaction. PULs were also impacted in nutrient limited conditions due to the presence of a bile acid. These PULs targeted a variety of substrates ranging from starches to host glycans. DCA, an inflammatory bile acid [61], impacted 14 of 21 differentially expressed PULS including ones that target host glycans and mucins. This suggest *B. theta* may be taking advantage of the inflammatory nature of this bile acid.

The native environment of *B. theta* is the mammalian gut, an environment that has a diverse pool of bile acids and as such it is not only coming into contact with one single bile acid at a time. To address this limitation and determine if the presence of these genes is required for colonization or creates a competitive advantage in a complex gut environment, we first used a gnotobiotic mouse model. In mono-colonization experiments, there were no significant differences past three days post challenge. This is consistent with work which showed no significant difference between CFUs in WT vs triple KO strains of *B. theta* in the feces after two weeks [8]. While slight differences in bacterial load were observed one day after challenge in the monocolonized mice, this difference was not recapitulated in the competition model. To determine if these enzymes impact colonization alongside a more complex microbiota this experiment was repeated using mice treated with cefoperazone to open up a niche for *B. theta* to colonize. Again, while we observed slight advantages in the WT over the triple KO strain, bile acid altering enzymes did not allow one strain to overwhelm the other.

Changes to the bile acid pool are associated with a wide variety of diseases including *C. difficile* infection (CDI) [10], Inflammatory bowel disease [11], metabolic syndrome [12], diabetes [25], hepatic fibrosis [24], different cancers [12, 13], and more recently depression [62]. Research on the enzymes that modify the bile acid pool is expanding with the recent finding that BSHs are associated with reconjugation of bile acids or MCBAs [15, 16, 33]. While *B. theta* was associated with the production of MCBAs [33], the specific enzyme that carries out this function has not been determined. Defining the mechanisms by which bacteria modify the bile acid pool will aid in our understanding of how to rationally manipulate the bile acid pool and the microbiota to treat many GI diseases.

## Data availability

## Acknowledgements

The authors thank the Roy J. Carver Biotechnology Institute at The University of Illinois for their assistance with RNA Sequencing. We thank J. Alfredo Blakeley-Ruiz for his assistance in annotating PULs in RNASeq analysis. We thank International Flavors & Fragrances Inc. for the funding they supplied for this work. This work was supported by the Molecular Biotechnology Training Program at NCSU (NIH NCSU MBTP T32 GM133366). CMT is a consultant for Ferring Pharmaceuticals and on the Scientific Advisory Board for Ancilia Biosciences. Matthew Foley is a consultant for Ancilia Biosciences.

## Supplemental Figures

**Supplemental Figure 1. Genes encoding bile acid altering enzymes impact both MIC and membrane integrity of *B. theta*.**

*B. theta* WT (A) and triple KO (B) strains were used to determine MICs in different bile acids. Bars represent mean MICs from (n = 2-3 biological replicates, n=2 technical replicates). Error bars represent standard error. Asterisks denote significance (p<0.0001) between conjugated and deconjugated by two way ANOVA with Tukey’s multiple comparisons test. *B. theta* WT (C) and triple KO (D) strains were incubated with different bile acids and stained with propidium iodide. Asterisks denote significant by two-way ANOVA with Dunnet’s multiple comparison test (*p<0.05;** p<0.01; **** p<0.0001) differences between the condition and the no bile acid or no BA control. (E) WT, *ΔbshA, ΔbshB, ΔhsdhA,* and the triple KO strains were incubated with CDCA and stained with propidium iodide. Asterisks denote significant (* p<0.05; *** p<0.0005) differences between each strain by two-way ANOVA with Tukey’s multiple comparison test.

**Supplemental Figure 2. Growth curves of WT *B. theta* in different bile acids with varying concentrations.**

WT and triple KO strains of *B. theta* were grown for 24 hr in TYG supplemented with different bile acids in varying concentrations. Error bars represent SD from n=3 biological replicates, n=2 technical replicates.

**Supplemental Figure 3. Genes encoding bile acid altering enzymes have minimal effects on *B. theta*’s fitness at 12 hours.**

WT and knockout strains of *B. theta* were grown for 12 hr in TYG supplemented with different bile acids. Error bars represent SD from n=2 biological replicates, n= 2 technical replicates. Dashed grey line represents the approximate starting CFU/mL added at 0 hr. Asterisks denote significant (p<0.05, p<0.01) differences from WT by one-way ANOVA with Dunnet’s multiple comparison test.

**Supplemental Figure 4. Isolation of purification of HSDH and BSHa.**

SDS PAGE gel (4-20%) of purified His-tagged BT_1911 (27.9 kDa) and BT_1259 (36.6 kDa) stained with Coomassie blue for visualization. 25 μg, 50 μg, and 100 μg of each protein was run alongside ladder labelled in kDa.

**Supplemental Figure 5. *hsdhA* is highly expressed in some bile acid conditions during mid-log phase of growth.**

Bars represent fold change in gene expression for genes encoding bile acid altering enzymes in different bile acids compared to expression in the absence of bile acid. Data was normalized using the ΔΔCt method using 16S rRNA as a housekeeping gene. Error bars represent standard error between replicates (n=3 biological, 3 technical replicates) each. Asterisks denote significant (* p<0.05; ** p<0.01; **** p<0.0001) differences between genes in each bile acid. by 2-way ANOVA with Tukey’s multiple comparison test.

**Supplemental Figure 6. Bile acids differentially impact the *B. theta* transcriptome in response to nutrient limiting conditions.**

Volcano plot of differentially expressed genes classified as a PUL or by KEGG orthology where p<0.05 by Wald test with Bonferroni correction and log_2_fold change > 2 or < −2 are plotted with a color. Number of genes increased in expression (top right corner) and decreased in expression (top left corner) are listed.

**Supplemental Figure 7. Bile acids impact multiple *B. theta* metabolic pathways in nutrient limited conditions.**

Heatmap of significantly differentially expressed genes with KEGG annotations where p<0.05 by Wald test with Bonferroni correction and log_2_fold change > 2 or < −2. Values represent the log_2_ fold change of genes that were significantly differentially expressed in nutrient limited conditions due to the presence of a bile acid.

**Supplemental Figure 8. Bile acid exposure impacts *B. theta* PUL expression in nutrient limited conditions.**

Heatmap of significantly differentially expressed genes annotated within CAZy as being part of a PUL where p<0.05 by Wald test with Bonferroni correction and log_2_fold change > 2 or < −2. Values represent the log_2_ fold change of genes that were significantly differentially expressed in nutrient limited conditions due to the presence of a bile acid.

**Supplemental Figure 9. *B. theta* taurine and glycine metabolism gene expression does not change in the presence of different bile acids in nutrient limiting conditions.**

Genes annotated in KEGG to be involved in taurine or glycine metabolism are listed. Values represent the log_2_ fold change in response to the addition of different bile acids in rich vs minimal cultures. No genes listed were found to be significant by Wald test with Bonferroni correction in any condition.

**Supplemental Table 1. Oligonucleotides used in this study.**

